# Decellularized Meniscus (MEND) as a biomaterial that supports stem cell invasion and chondrogenesis

**DOI:** 10.1101/2025.10.16.682874

**Authors:** Hannah M. Bonelli, Sophia E. Klessel, Cristina Barbella, Kyra W.Y. Smith, Riccardo Gottardi

**Author notes:** Corresponding author: Riccardo Gottardi, PhD, Children’s Hospital of Philadelphia, 3615 Civic Center Blvd, Philadelphia, PA 19104, USA.

## Abstract

**BACKGROUND:** Cartilage damage affects 25 million people globally each year. Tissue engineering strategies such as microfracture and matrix induced autologous chondrocyte implantation (MACI) are currently being used in the clinic; however, they are accompanied by their own limitations such as donor site morbidity, rapid clearance from the injury site, and extensive cost. To overcome these limitations, the tissue engineering field has shown increasing interest in the use of decellularized extracellular matrix (dECM) biomaterials due to their heightened integration with native tissue and regeneration rates.

**METHODS:** The Gottardi Lab has developed a new dECM material sourced from porcine <u>men</u>iscus <u>d</u>ecellularization (MEND), in which elastin fibers are removed via enzymatic digestion, resulting in channels that can be easily recellularized.

**RESULTS:** In this work we demonstrate that MEND can be seeded with bone-marrow derived mesenchymal stem cells (MSCs), achieving a uniform distribution of cell nuclei throughout the cross section of the scaffold. We also show that MEND retains its native structure in the presence of MSCs and can support chondrogenesis comparably to other commonly used tissue engineering materials such as methacrylated type I collagen and gelatin/hyaluronic acid hydrogels.

**CONCLUSION:** Overall, MEND is a promising new dECM biomaterial for cartilage regeneration.

## 1 Introduction

Cartilage damage affects 25 million people annually [1]. As a direct effect, about 200-300k cartilage procedures are performed just on knees each year in the U.S. alone [2]. These cartilage injuries can be caused by a variety of occurrences such as trauma, sports injuries, and wear and tear overtime [3]. This leads to joint pain, clicking, stiffness, inflammation, and an overall decreased quality of life [4]. Palliative care options such as nonsteroidal anti-inflammatory drugs and physical therapy can be used to alleviate pain but they merely treat symptoms rather than repair the cartilage damage [4]. Ultimately, total joint replacement is an end-stage treatment after the exhaustion of palliative care options but this is extremely invasive, presents potential infection risk after major open surgery, and implant life spans are still limited [5]. Instead, regenerative medicine and tissue engineered solutions may overcome these limitations or at least postpone the need for total joint replacement and have in fact been in practice for decades. Microfracture is the most common tissue engineering procedure, however, it results in fibrocartilage which has limited durability in the joint compared to the hyaline articular cartilage [6]. Another more involved approach, matrix induced autologous chondrocyte implantation (MACI) is an extremely effective solution. However, MACI requires two procedures which increases cost and recovery time and decreases rates of patient compliance [7,8]. Furthermore, it is only effective for an isolated and relatively small cartilage defect which excludes the majority of patients who present more advanced cartilage damage. To overcome some of these limitations, additional tissue engineering approaches have been explored for cartilage repair and regeneration such as synthetic polymers, hydrogels, and decellularized extracellular matrix. All are capable of presenting translatable solutions that can support chondrogenesis.

Synthetic polymers have been used as scaffolds for orthopedic engineered constructs with strong mechanics, tunable porosity, and potential for functionalization [9]. However, the synthetic nature of the materials may result in poor integration, risk of foreign body response, and poor cell adhesion [10]. Oftentimes they require a pre-coating with cell adhesion elements such as serum or collagen, or functionalization with integrin binding motifs [10]. Furthermore, synthetic materials often present challenging degradation rates that may not match new tissue regeneration. This may cause a potential reaction of host tissues to scaffold degradation products and in some cases, scaffolds may fail to degrade at all [11]. Overall, synthetic materials possess great potential in terms of tunability but are often accompanied by equally challenging limitations that hinder their translatability.

To overcome these limitations, hydrogel-based biomaterials derived from natural sources such as collagen, gelatin, and hyaluronic acid have been widely explored, presenting good integration potential with surrounding tissues and robust cellular adhesion [12]. Bosnakovski et al. studied type I collagen hydrogels for chondrogenesis and although MSC mediated matrix secretion occurred, it resembled that of fibrocartilage, missing key elements of the desired hyaline cartilage phenotype [13]. Additionally, Galois et al. found that collagen hydrogels lacked initial mechanical properties and tended to contract in the presence of cells, making it difficult to study tissue regeneration due to non-uniform matrix secretion [14]. Gelatin and hyaluronic acid hydrogels have been comparatively more successful in promoting hyaline cartilage matrix that is rich in collagen II. However, these hydrogels often require high cell densities to reach desired matrix production, in part because the abundant crosslinking that provides stability to the hydrogel may be difficult for cells to remodel, ultimately leading to limited translatability [15]. Overall, hydrogels are a promising biomaterial for cartilage regeneration but often result in uneven matrix formation and frequently present initial mechanical properties that are insufficient for translation without extensive *in vitro* pre-culture.

Decellularized extracellular matrix-based (dECM) biomaterials present an exciting option as they have been shown to possess increased integration and support for regeneration [16]. dECM scaffolds have been fabricated from a wide variety of tissues and have also been used for many applications. For instance, porcine urinary blader has been used as a dECM scaffold to engineer the temporomandibular joint disc, demonstrating cellular infiltration *in vivo* [17]. Likewise, Sladkova et al. found that dECM scaffolds derived from bovine bone could support calcified bone formation and cell viability just as well as human dECM bone scaffolds [18]. Given the high density and avascular nature of cartilage tissue, dECM scaffolds derived from cartilage often face additional obstacles. Zeng et al. created dECM scaffolds from porcine articular cartilage; however, they opted to pulverize the tissue, forming an injectable hydrogel slurry [19]. While it was evident that cartilage regeneration occurred, they lost the natural structure and mechanical properties of the articular cartilage in order to achieve cellular encapsulation. In our previous work, we have faced similar challenges and resorted to combining dECM derived from tendon and cartilage with methacrylated gelatin. We created a dECM/hydrogel system that contained biological cues for cell specification and hydrogel components for stability, however initial mechanical properties did not match those of native tissue [20]. Nürnberger et al. developed a CO_2_ lasered scaffold that consisted of precisely drilled decellularized articular cartilage to create a scaffold with controlled channels [21]. Although this technique was successful in overcoming previously mentioned limitations such as maintaining native structure and achieving cell migration, cells remained nonuniform after initial seeding and required significant matrix remodeling to achieve uniform matrix production across scales. In the Gottardi Lab, we have developed a new scaffold based on porcine <u>men</u>iscus <u>d</u>ecellularization (MEND) [22,23]. MEND is fabricated from the red-red zone of porcine meniscal tissue, consisting of fibrocartilage that is rich in type I collagen, hyaluronic acid, blood vessels, and elastin fibers. MEND is obtained from the selective removal of elastin without otherwise impacting meniscal structure, which results in a wealth of channels that cells can infiltrate for re-cellularization.

In this study, we seeded MEND with MSCs to compare the chondrogenic response against that of MSCs in 3D hydrogels with controlled composition. Since MEND is primarily composed of type I collagen, we chose to compare MSC differentiation in MEND to that of MSCs in methacrylated type I collagen (ColMA) hydrogels. Similarly, hyaluronic acid is also a component of MEND matrix, so we opted to include a methacrylated gelatin-hyaluronic acid (GelMA/HAMA) hydrogel as well. Given that GelMA/HAMA hydrogels and MEND are composed of matrix components typically produced by MSCs during differentiation, we used fluorescent non-canonical amino acid tagging (FUNCAT) and click chemistry to fluorescently label cell secreted neo-matrix and distinguish it from that of the scaffold. By comparing chondrogenesis of MSCs in MEND with ColMA and GelMA/HAMA hydrogels, we assessed MEND’s ability to support chondrogenesis (chondroconductivity) and whether the meniscal ECM has any specific pro-chondrogenic effect on MSCs (chondroinductivity) *in vitro*.

## 2 Methods

### 2.1 Cell Sources

This study was conducted using human bone marrow mesenchymal stem cells extracted from the iliac crest (RoosterBio; Frederick, MD) at passage 5. A total of 5 donors were used to account for the donor variability of using human sourced cells. Donors were 4 male and 1 female, ranging from 19 to 29 years of age. For each donor, cells were expanded in RoosterBasal medium (RoosterBio; Frederick, MD) and lifted with trypLE Express (ThermoFisher Scientific; Waltham, MA) at 80-100% confluency. Cells were transferred in DMEM (ThermoFisher Scientific; Waltham, MA) containing 10% fetal bovine serum (FBS, Avantor VWR; Radnor, PA) and 2% penicillin, streptomycin, and fungizone (PSF, ThermoFisher Scientific; Waltham, MA). For MEND seeding, cells were resuspended in DMEM containing 1% FBS and 2% PSF. Constructs underwent chondrogenesis for 21 days in chondrogenic medium (DMEM, 2% PSF, 10μg/mL Insulin-Transferrin-Selenium, 40μg/mL L-proline, 0.1 µM Dexamethasone, 50 µg/mL Ascorbic Acid and 10ng/mL Transforming Growth Factor-β3 (TGF-β3)). Constructs undergoing chondrogenic FUNCAT culture received chondrogenic medium made with methionine free DMEM supplemented with 1 µL/mL of Click-IT™ L-Homopropargylglycine (HPG, ThermoFisher Scientific; Waltham, MA) and L-cystine. Medium was renewed every other day for the duration of culture. All constructs were cultured on a shaker placed within an incubator at 37C shaking at 100 RPM except the pellets which were cultured in static conditions.

### 2.2 Pellet preparation

Cells were resuspended at a density of 200k cells/200 µL of media. 200 µL of cell solution was added to each well of a U-bottom 96-well plate and then the plate was centrifuged at 300XG for 10 minutes. The plate was incubated at 37C for 24 hours (24h) and then switched to chondrogenic culture.

### 2.3 Collagen hydrogel fabrication and seeding

Lyophilized bovine derived methacrylated type I collagen (ColMA, Advanced BioMatrix; Carlsbad, CA) was resuspended in 20 mM acetic acid according to the manufacturer instructions. Briefly, solution rotated at 4C for 2-3 days to allow ample mixing. During each hydrogel fabrication, 3% 1M NaOH, 10% 10X phosphate buffered saline (PBS), and 2% lithium phenyl-2,4,6-trimethylbenzoylphosphinate (LAP) were added for a final collagen concentration of 6 mg/mL.

Cell solution was added to the hydrogel mixture so that the final construct density was 10 million cells/mL. Solution was pipetted into a sterile 80 µL silicone mold, thermo-crosslinked for 30 minutes at 37C and then photo-crosslinked under UV light for 2.5 minutes. Constructs were cultured for one day in DMEM with 10% FBS and 2% PSF prior to switching to chondrogenic medium.

### 2.4 Gelatin/Hyaluronic acid hydrogel fabrication and seeding

Lyophilized porcine derived methacrylated gelatin (GelMA, Advanced BioMatrix; Carlsbad, CA) and hyaluronic acid (HAMA, Advanced Biomatrix; Carlsbad, CA) were separately resuspended in 1.5 mg/mL solution of LAP and 1X PBS and placed on a 37C shaker overnight. Then, GelMA was added to HAMA to a final concentration of 5% GelMA, 2.5% HAMA, 0.15% LAP and mixed again overnight.

Cells were seeded at a final construct density of 10 million cells/mL and pipetted into sterile 20 µL silicone molds, then photo-crosslinked under UV for 1-2 minutes on each side. Hydrogels were moved to a U-bottom 96-well plate and then were switched to chondrogenic culture after 24h.

### 2.5 MEND fabrication and seeding

Adult porcine menisci (Collagen Solutions) were sectioned radially with a thickness of 0.5-1 mm (Figure 1A-D). Menisci sections underwent two freeze/thaw cycles at 1h each. Then sections underwent a third freeze/thaw cycle in 10 mM Tris-base (pH 8). Samples were then treated with 0.1% pepsin in 0.5M acetic acid for 24h at 37C while shaking in an Erlenmeyer flask (Figure 1E), followed by a 24h wash in 1X PBS and an incubation with an elastase solution (0.3 U/mL elastase, 0.2M Tris-base pH 8.6) for 24h at 37C under shaking at 180 RPM (Figure 1F). Sections were then washed in PBS for 24h to confirm decellularization (Figure 1G) and a 6 mm biopsy punch was used to create cylindrical scaffolds from the red-red zone of the meniscal cross section (Figure 1H). These scaffolds were washed for 24h at 37C in DMEM supplemented with 20% FBS and 2% PSF.

**Figure 1.**
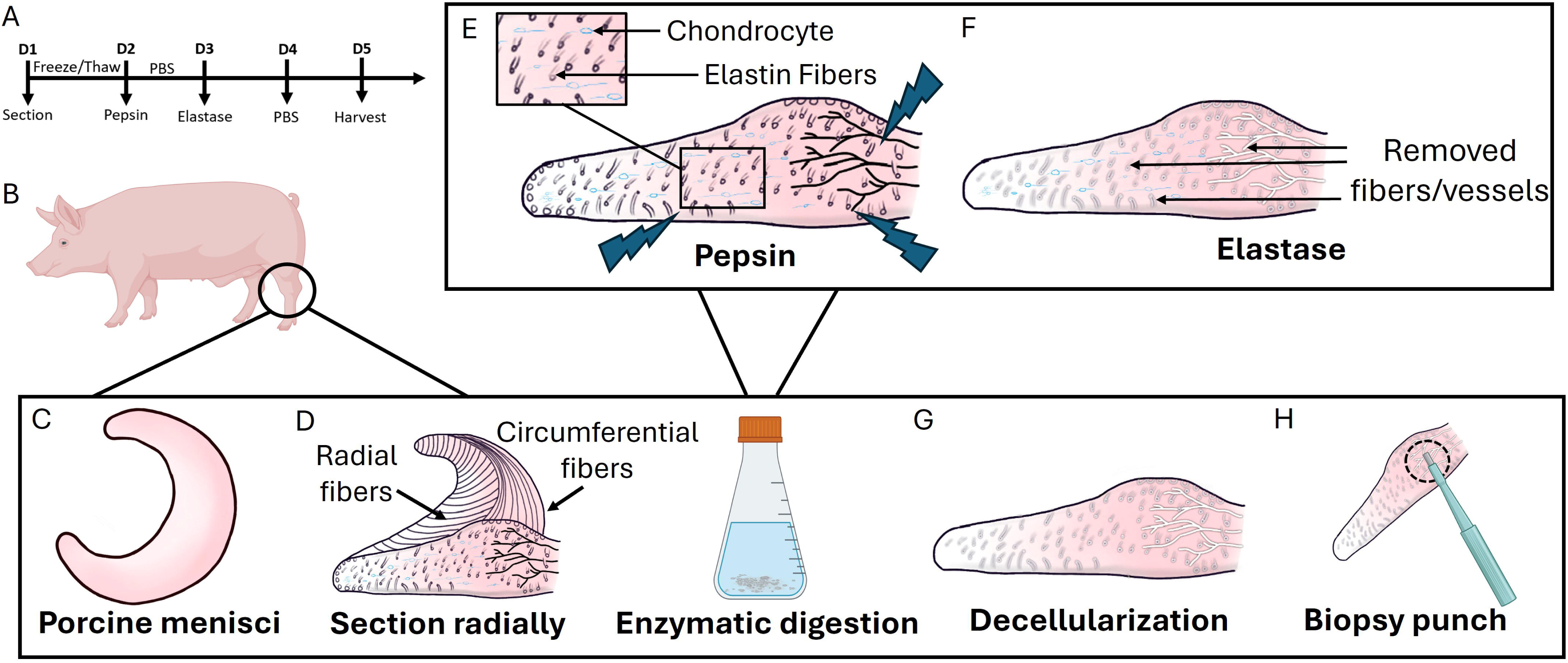
Decellularized meniscus (MEND) fabrication. **(A)** Timeline of the fabrication process. MEND undergoes 3 freeze/thaw cycles, then is subjected to 24 hours in pepsin-acetic acid solution and 24 hours in elastase. MEND is washed in 1X PBS for 24 hours and stored at - 20C until use. **(B and C)** Menisci are acquired from adult Yorkshire pigs and are **(D)** sectioned radially to a thickness of 0.5-1 mm. Then, sections are subjected to a **(E)** pepsin and acetic acid solution to loosen the tissue and **(F)** elastase solution to remove the circumferential elastin fibers. **(G)** At this point, the biomaterial is decellularized and **(H)** ready to be biopsy punched to its desired size. For this study, we used a 6 mm biopsy punch.

MEND was seeded using a transwell plate (Corning Inc.; Corning, NY) and FBS serum gradient. First, sterile 6.5 mm silicone rings were placed in the top insert of the transwell. MEND was washed 3 times in DMEM containing 1% FBS and then tightly fitted inside the silicone ring. Cell were seeded on top of MEND at 100k, 400k, or 600k cells/construct. DMEM with 20% FBS and 2% PSF was added to the bottom of the transwell, creating a serum gradient to drive cell migration into the construct. Media was renewed every other day for 5 days, then constructs were transferred to a 12 well plate containing chondrogenic medium for 21-day culture.

### 2.6 Unconfined compression testing

Native meniscus and MEND underwent unconfined compression testing using a custom designed computer-controlled testing apparatus [24]. We used a ramp velocity of 10 µm/s until reaching 10% strain. All constructs had a thickness of ∼0.5-1 mm and a diameter of ∼6 mm.

### 2.7 Histology and Immunofluorescence

All conditions were collected at Day 0 and Day 21 except pellets, which are not yet formed at Day 0. Samples were washed 3 times in 1X PBS, fixed in 10% formalin for 1h, then washed 3 times in 1X PBS. Pellet, ColMA, and MEND samples underwent sequential ethanol and xylene dehydration, and then were embedded in paraffin and sectioned at 8 µm thickness using a microtome (Mircom HM 355S, ThermoFisher Scientific; Waltham, MA). Due to high water content and to avoid swelling after sectioning, GelMA/HAMA samples underwent a sucrose gradient (10%, 20%, 30% in 1X PBS) prior to embedding in optimal cutting temperature compound (OCT, Sigma-Aldrich; St. Louis, MO) using dry ice and isopentane (ThermoFisher Scientific; Waltham, MA). Samples were sectioned at 8 µm thickness with a cryotome (Microm HM 505E, GMI, Ramsey, MN).

After deparaffinization and rehydration, sections were stained for histology and immunofluorescence. To minimize binding of non-sulfated glycosaminoglycans (GAGs) that are part of the GelMA/HAMA hydrogel itself, samples were first incubated with hyaluronidase for 1h at 37C and then washed 3 times with 1X PBS. Alcian Blue staining was conducted using a pH of 1.0 for sulfated GAG detection only. After 30 minutes of Alcian Blue, slides were washed using 0.1M hydrochloric acid (HCl) to prevent an increase in pH which could lead to further binding. 3 slides per sample were stained and analyzed. Slides were imaged at 20X magnification using an inverted microscope (BZX-810 Keyence, Blue Bell, PA).

All FUNCAT cultured samples were cryo-sectioned as described above and cryo-preserved to maintain fluorescent capabilities. Day 21 ColMA, GelMA/HAMA, and MEND samples were stained for immunofluorescence imaging using a biotinylated hyaluronic acid binding peptide (AMSBio; Cambridge, MA). Slides were incubated in blocking buffer (1X PBS, 0.1% Triton-X, 2% bovine serum albumin) for 2h at room temperature and then incubated with the hyaluronic acid binding peptide overnight at room temperature. Slides were then washed 3 times in 1X PBS and incubated with Streptavidin Alexa647 for 1h at room temperature. After washing 3 times in 1X PBS, FUNCAT staining was performed according to the manufacturer instructions. Briefly, slides were washed twice in 2% BSA in 1X PBS and then incubated for 20 minutes in 0.5% Triton X-100. Slides were washed again in 3% BSA solution and incubated with Click-iT® reaction cocktail (ThermoFisher Scientific; Waltham, MA) at room temperature for 30 minutes. Slides were then washed in Click-iT® rinse buffer (ThermoFisher Scientific; Waltham, MA) and incubated for 30 minutes in NuclearMask™ Blue Stain (1:2000 in PBS, ThermoFisher Scientific; Waltham, MA). Lastly, slides were washed twice with 1X PBS and coverslips were mounted with Fluoromount. Slides were then imaged with an inverted fluorescent microscope (BZX-810 Keyence, Blue Bell, PA).

### 2.8 Biochemical analysis

Samples were collected at both Day 0 and Day 21 timepoints for all conditions. For ColMA, one construct per donor per condition was sufficient for data collection. GelMA/HAMA required pooling of 3 constructs, while pellets required pooling of 5. MEND biochemical analysis was carried out using half of a construct. After washing 3 times in PBS, samples were lyophilized for 24h, dry weight was recorded, then samples were digested in 0.125 mg/mL papain (Sigma-Aldrich; St. Louis, MO) in PBS with 5mM ethylenediaminetetraacetic acid (EDTA, Sigma-Aldrich; St. Louis, MO) and 5mM L-cysteine (Sigma-Aldrich; St. Louis, MO) at 60C for 24h. GelMA/HAMA were first incubated in 5 mg/mL hyaluronidase (Sigma-Aldrich; St. Louis, MO) at 37C for 24h, prior to the same papain digestion. When samples were completely digested with no noticeable precipitate remaining, the solution was removed from the oven and centrifuged at 13000XG for 5 minutes and the supernatant was used for biochemical assays.

For total DNA quantification, Quant-iT™ PicoGreen™ (ThermoFisher Scientific; Waltham, MA) was conducted according to manufacturer protocol. Briefly, samples were diluted 1:10 in 1X TE buffer and 100 µL was added to each well of a black 96-well plate in duplicate. PicoGreen™ was diluted 1:200 in 1X TE and 100 µL was added to each well of the plate. The plate was incubated for 5 minutes at room temperature, protected from light. Standard curve was established using lambda DNA in digestion buffer. Standard was diluted in 1X TE buffer to maintain consistency between the standard and the samples. The plate was read at 480 nm excitation and 520 nm emission using a plate reader (BioTek Synergy H1, Santa Clara, CA).

For total GAG quantification, supernatant of all conditions was diluted 1:5 in deionized water. To make the dimethylmethylene blue (DMMB) solution, 1 g of sodium formate was dissolved in deionized water, and then 1 mL of formic acid was added to maintain a pH of 3.5 of lower. A solution of 8 g DMMB in 2.5 mL of ethanol was added, followed by extensive vortexing and sonication to properly dissolve the DMMB dye. Chondroitin-6-sulfate diluted in deionized water was used to establish the standard curve. 40 µL of standard solution or diluted sample solution was added to each well of a clear 96-well plate in duplicate. Then, 250 µL of DMMB solution was added to each well and the plate was read immediately at both 540 and 595 nm.

### 2.9 Real-time quantitative reverse transcription polymerase chain reaction (RT-qPCR)

Samples were collected at both Day 0 and Day 21 timepoints for all conditions. For ColMA, one construct per donor per timepoint was sufficient for data collection. GelMA/HAMA and MEND required pooling of 3 constructs, while pellets required pooling of 5.

For RNA extractions, samples underwent 3 free/thaw cycles in liquid nitrogen and were then pulverized with a micro-homogenizer and pestle while submerged in Trizol (Thermo Fisher Scientific, Waltham, MA). The top aqueous layer was extracted from the tube and then samples underwent manufacturer protocol according to the RNeasy Plus Mini Kit (Qiagen; Hilden, Germany).

RT-qPCR was conducted using SYBR Green (Applied Biosystems; Waltham, MA). See Table 1 for forward and reverse primer sequences (Integrated DNA Technologies; Coralville, IA). The RT-qPCR plate ran on a QuantStudio 7 and data were reported as fold change of relative gene expression (2^-ΔΔct^).

**Table 1.**
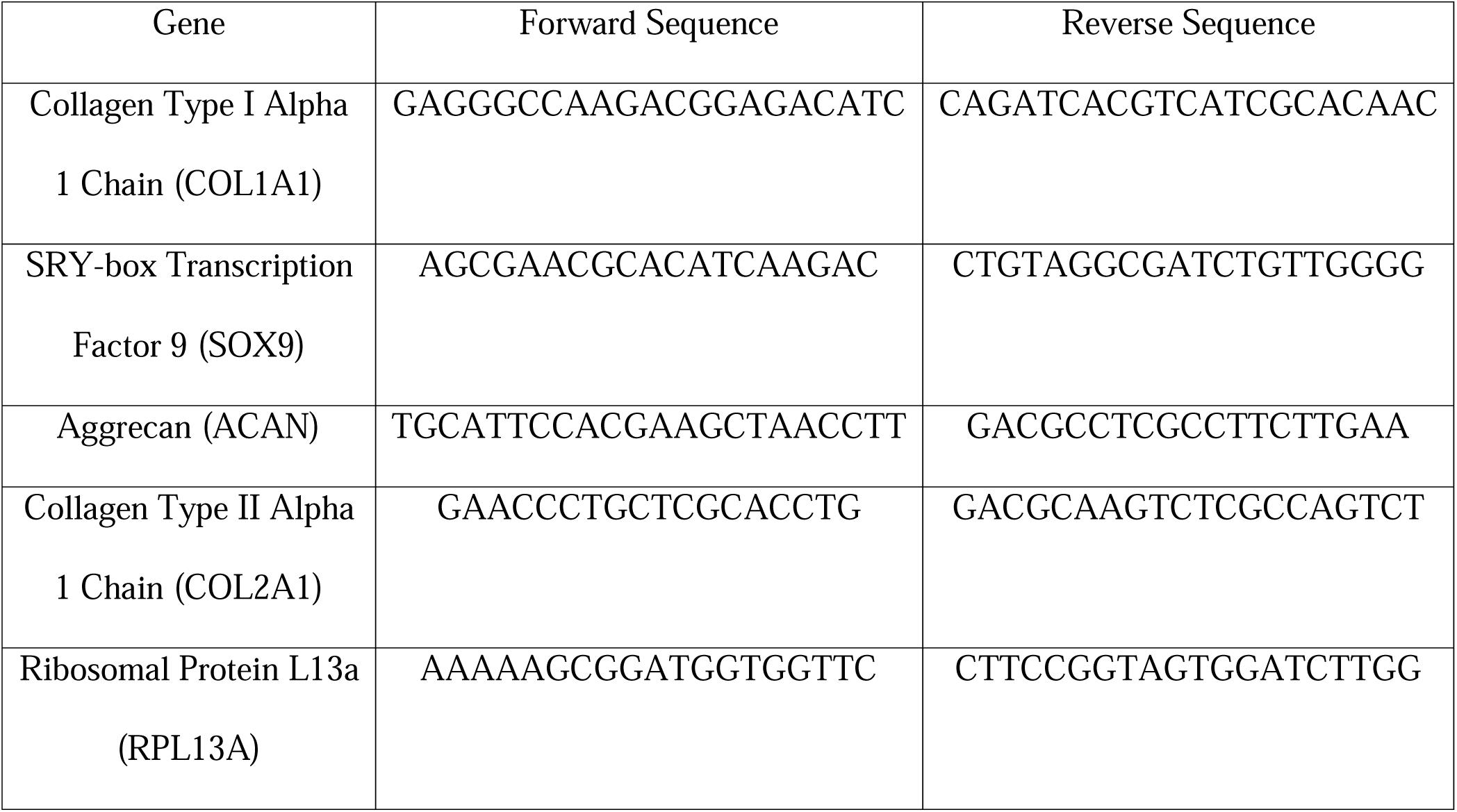
Forward and reverse sequences for RT-qPCR primers.

### 2.10 Statistical analysis

All statistical analyses were conducted using GraphPad Prism 7. A 1- or 2-way ANOVA followed by Tukey’s *post hoc* test was used when statistical measures included one or two independent variables, respectively. A student T-test was conducted for comparison of two groups. In all graphs, *p<0.05, **p<0.01, ***p<0.001, and ****p<0.0001.

## 3 Results

### 3.1 MEND fabrication creates channels

We fabricated MEND by selective enzymatic digestion of elastin bundles as previously described (Figure 1) [22]. MEND’s structure was compared to native porcine meniscus using hematoxylin and eosin (H&E) staining, demonstrating increased porosity and vertically oriented channels throughout the cross section of the material (Figure 2A and B). This resulted in a 4-fold increase in porosity in MEND (Figure 2C). Moreover, Modified Hart’s staining confirmed the removal of elastin in MEND, marked by the lack of purple hue compared to native (Figure 2D and E). Notably, after the removal of elastin fibers we observed a 2-fold decrease in compressive modulus (Figure 2F). DAPI staining conducted on representative sections from multiple planes within MEND and native meniscus showed the removal of cell nuclei (Figure 2G and H). This was further confirmed by DNA quantification using PicoGreen biochemical analysis, indicating that MEND is decellularized below the FDA standard of 50 ng/mg of tissue (Figure 2I). Alcian blue staining was performed to assess the impact of the fabrication process on GAG retention (Figure 2J-K). Biochemical analysis showed a significant decrease of GAGs in MEND (Figure 2L), dropping from 18.7 µg/mg of tissue in native to 0.57 µg/mg of tissue in MEND. Despite this loss, MEND retains much of its native structure with uniform channels throughout; an ideal characteristic for cell seeding.

**Figure 2.**
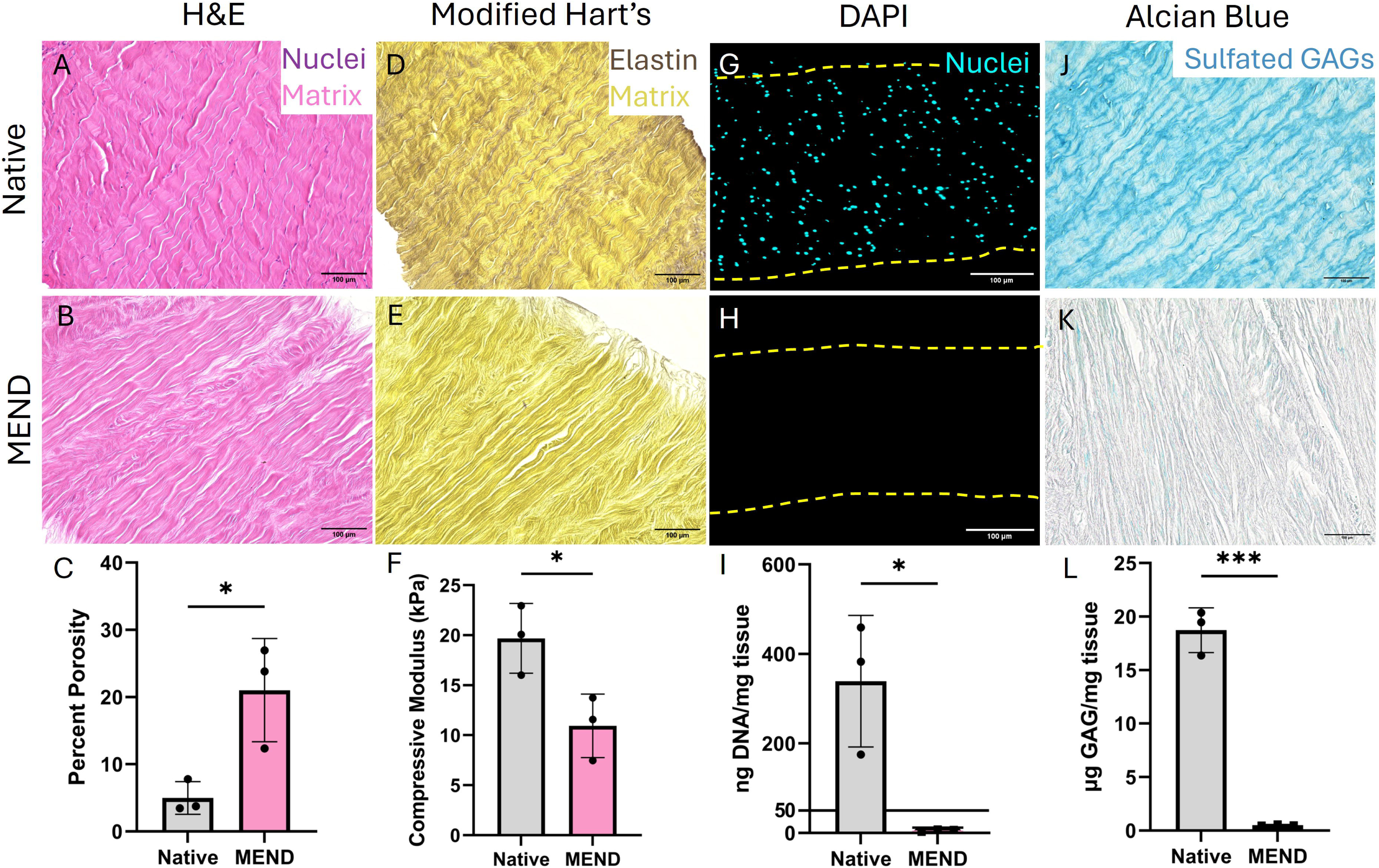
MEND fabrication creates channels. Hematoxylin & eosin (H&E) of **(A)** native porcine meniscus and **(B)** MEND demonstrating cell nuclei in purple and matrix in pink. **(C)** Percent porosity increased significantly between native meniscus and MEND. Modified Hart’s of **(D)** native meniscus and **(E)** MEND shows a decrease in elastin (purple) and the formation of channels in MEND. **(F)** Compressive modulus shows a decrease in value between native meniscus and MEND. DAPI staining of **(G)** native meniscus and **(H)** MEND shows a decrease in cell nuclei. **(I)** Biochemical analysis of DNA content using PicoGreen confirms a significant decrease in cell presence below the FDA standard of 50 ng/mg tissue. Alcian Blue of **(J)** native meniscus and **(K)** MEND shows a decrease in sulfated glycosaminoglycan (GAG) content and **(L)** biochemical analysis for quantitative GAG content confirms this decrease. A, B, D, E, J, and K: Scalebar = 100 µm; 20X magnification, G and H: Scalebar = 100 µm; 10X magnification. Student T test, *p<0.05, ***p<0.001

### 3.2 MEND structure allows for uniform stem cell seeding

MEND seeding followed a 6-day time course (Figure 3A), where bone marrow derived MSCs were seeded on top of MEND in a transwell plate with a serum gradient to drive chemotaxis through the scaffold (Figure 3B). Three different MSC densities were tested to identify the most efficient seeding. After 5 days, calcein AM (ThermoFisher Scientific; Waltham, MA) staining showed that the 100k seeded cells condition resulted in the most even distribution of cells on top of MEND (Figure 3C). Moreover, many cells were also visible below the surface of MEND, further indicating their migration into the scaffold (Figure 3D). This was confirmed with DAPI staining, which showed a uniform distribution of cells throughout the scaffold’s cross section (Figure 3E). Seeding of 400k cells also resulted in a consistent distribution on the top surface (Figure 3F), but we were unable to visually identify cells migrating through to the bottom of the scaffold (Figure 3G). In fact, DAPI displayed cell migration through the scaffold, however, not as efficiently as when seeded with 100k MSCs (Figure 3H). Finally, seeding 600k cells per MEND resulted in the formation of a cell sheet on top (Figure 3I). Although we observed cells on the bottom of the scaffold (Figure 3J), DAPI proved that cells primarily remained on the top with a non-uniform distribution throughout the cross section (Figure 3K). Overall, as cell seeding density increased above 100k cells, the cell migration into the scaffold decreased, suggesting that cells interact with each other on the surface instead of migrating towards the nutrient rich environment. Hence, we used the 100k cells per MEND condition throughout the rest of the study.

**Figure 3.**
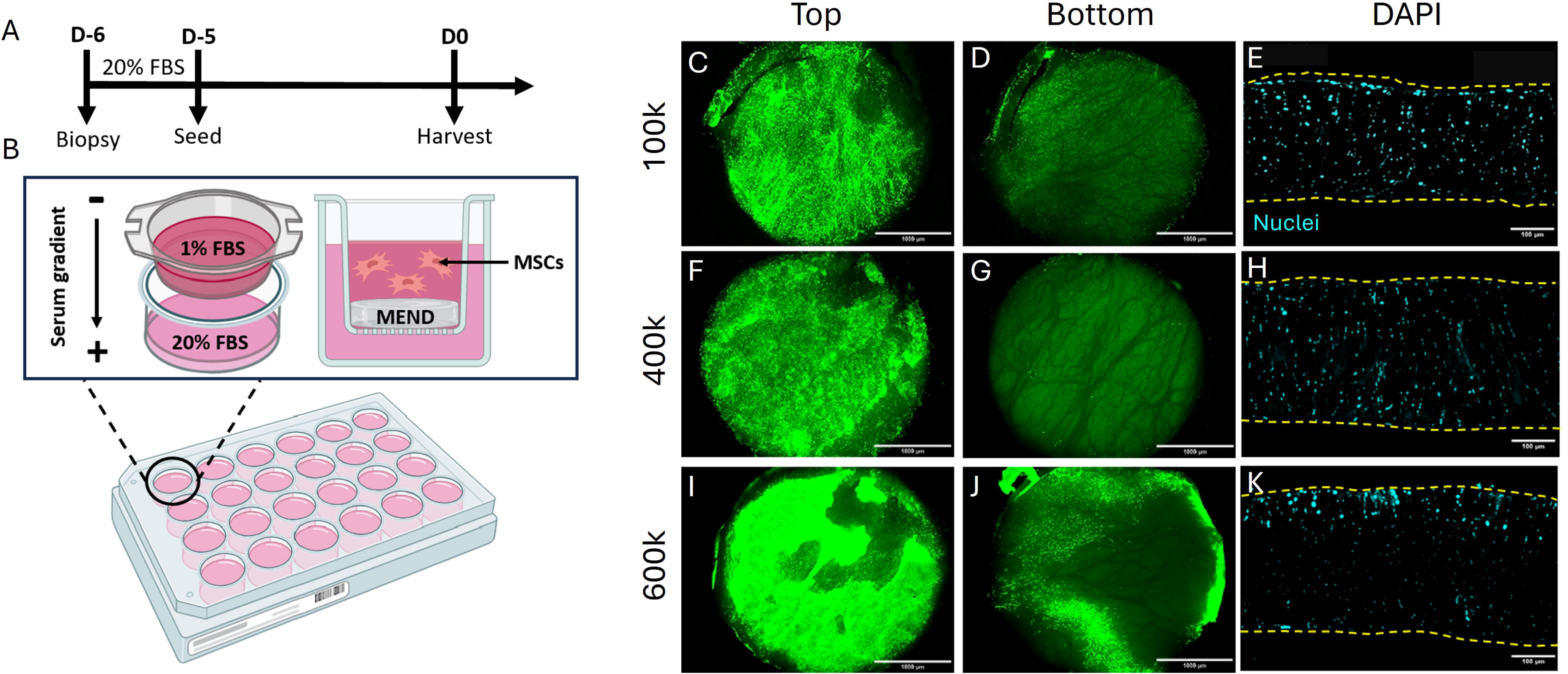
MEND structure allows for uniform stem cell seeding. **(A)** Timeline of MSC seeding on MEND. 6 days before harvest, MEND is washed with 20% fetal bovine serum (FBS) for 24h. MSCs are seeded on MEND and allowed time to migrate through for 5 days before being collected or advancing to chondrogenic differentiation. **(B)** Schematic of MEND MSC seeding. MEND was placed at the bottom of the transwell insert and MSCs suspended in media containing 1% FBS were seeded on top. Media containing 20% FBS was placed in the bottom to create a serum gradient to encourage cells to migrate through the scaffold and towards the nutrient-rich environment. **(C)** Top and **(D)** bottom images of Calcein AM stained MSCs seeded on MEND for 5 days at a density of 100k/MEND. **(E)** DAPI staining of 100k MSCs/MEND shows an evenly distributed array of cell nuclei over the cross section of the scaffold. **(F)** Top and **(G)** bottom images of Calcein AM stained MSCs seeded on MEND for 5 days at a density of 400k/MEND. **(H)** DAPI staining of 400k MSCs/MEND shows uniform distribution of cell nuclei throughout the cross section of MEND. **(I)** Top and **(J)** bottom images of Calcein AM stained MSCs seeded on MEND for 5 days at a density of 600k/MEND. **(K)** DAPI staining of 600k MSCs/MEND shows poor migration through the cross section of the scaffold, with the best migration only occurring at the top. Top and bottom images were taken at 2X magnification, scalebar = 1000 µm and DAPI at 10X magnification, Scalebar = 100 µm.

### 3.3 MEND supports chondrogenesis of MSCs

Real time quantitative reverse transcription polymerase chain reaction (RT-qPCR) was used to quantify the chondrogenic response of MSCs within each material condition, using MSC pellets as positive control. We compared the gene expression of *SRY-Box Transcription Factor 9 (SOX9,* Figure 4A), an early marker and master regulator of chondrogenesis, *aggrecan* (*ACAN*, Figure 4B), a late chondrogenic marker, and of C*ollagen Type I Alpha 1 Chain* (*COL1A1*) and *Collagen Type II Alpha 1 Chain* (*COL2A1*), which are prominent matrix components of fibrocartilage and hyaline cartilage, respectively (Figure 4C and D). *SOX9*, *ACAN*, and *COL2A1* were upregulated in MEND constructs and followed similar trends as the other materials tested. Although the high variability of biological replicates using human cells does not result in statistical significance, we can observe some trends. *SOX9* expression increased 2-fold between day 0 and day 21 for all conditions. *ACAN* expression between days 0 and 21 increased over 20-fold for all conditions. The expression levels for these genes suggest robust and mature chondrogenesis on day 21 and provides evidence that MEND is comparable to the other conditions in supporting expression of chondro-specific matrix components. *COL2A1* expression also increased between days 0 and 21 by 5-fold in all conditions. The same was true for *COL1A1* which is a common occurrence during *in vitro* chondrogenic culture using MSCs[25–27]. The ratio of *COL2A1*/*COL1A1* increased 2-fold by day 21 for the MSC pellets and GelMA/HAMA conditions, suggesting that newly deposited matrix resembled that of hyaline cartilage. ColMA did not show an increase in the ratio of *COL2A1*/*COL1A1* on day 21 compared to day 0, suggesting that ColMA may support a more fibrocartilage-like matrix in the presence of MSCs. MEND performed in between these outcomes, with no increase on day 21 compared to day 0. Even in this respect, MEND performs comparably to the other scaffold conditions and to the pellet control without altering the cellular response to differentiation medium (Figure 4E).

**Figure 4.**
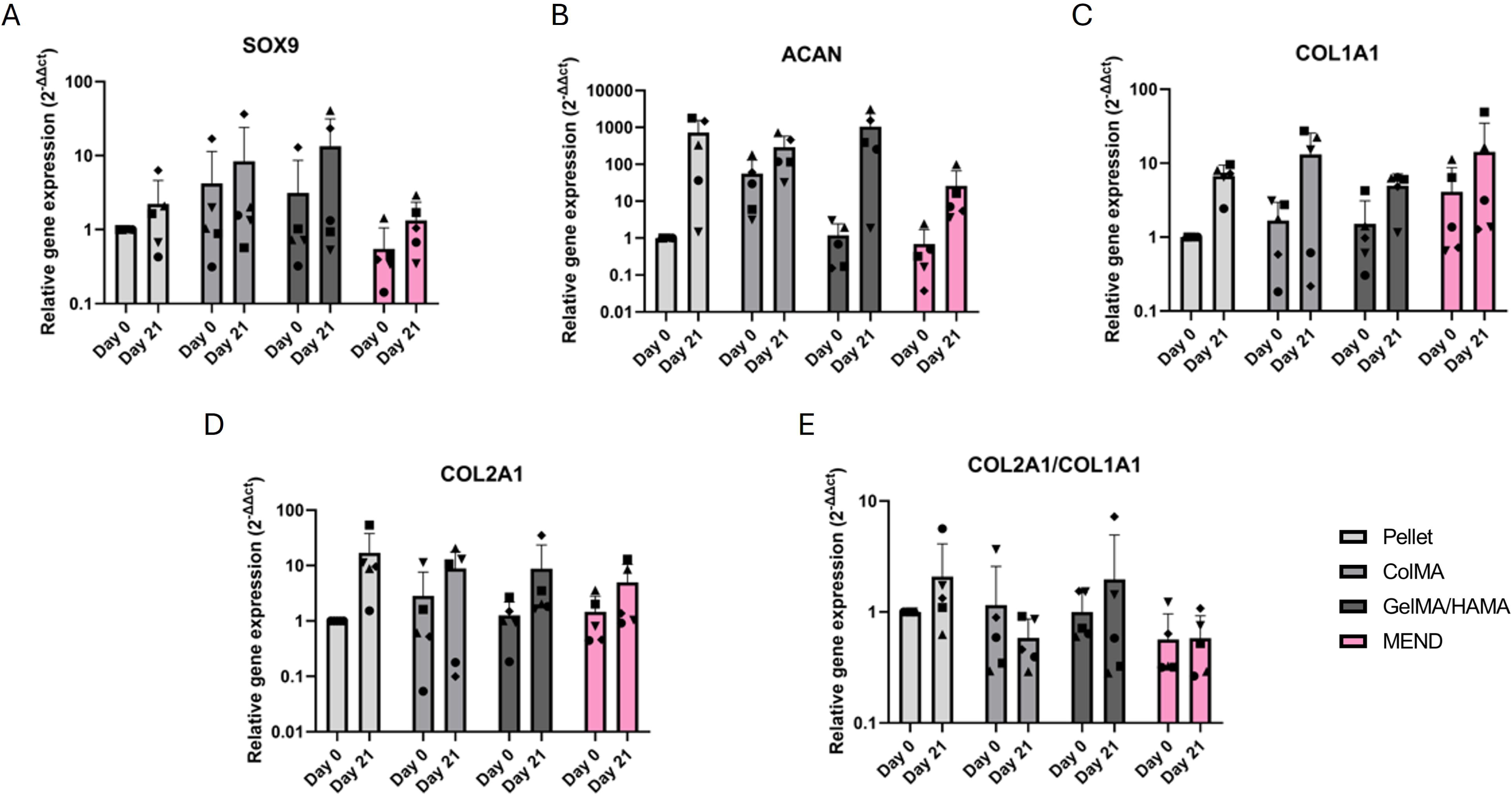
MEND supports chondrogenesis of MSCs. Real time-quantitative polymerase chain reaction (RT-qPCR) was conducted for **(A)** SOX9, **(B)** ACAN, **(C)** COL1A1, and **(D)** COL2A1 for all materials, including the chondrogenic gold standard pellet culture. Relative gene expression was normalized to RPL13A and day 21 samples were compared to day 0. **(E)** The ratio of COL2A1/COL1A1 for all materials as a representation of hyaline type expression. Each data point represents a human MSC biological replicate performed in technical duplicate. ● = donor 1, ▪ = donor 2, ▲ = donor 3, ▼ = donor 4, and ♦ = donor 5. A one-way ANOVA was conducted on all comparisons.

### 3.4 MEND supports uniform matrix secretion and higher GAG production per cell

The sulfated GAGs present in each construct were assessed via Alcian Blue at a pH of 1.0 for mucopolysaccharide specificity. All biomaterials in this study supported an increase in GAG secretion between day 0 and day 21. However, the newly produced matrix was non-uniformly distributed in both day 21 MSC pellets (Figure 5A) and day 21 ColMA gels (Figure 5B). Furthermore, the ColMA samples drastically changed their shape with substantive contraction. Differently, the GelMA/HAMA and MEND constructs presented uniform GAG distribution throughout the samples and maintained their original dimensions with no obvious contraction throughout the entire 21-day culture period (Figure 5C and D). However, GelMA/HAMA as well as MEND, albeit to lesser extent, already contain a base level of GAGs which lightly stains by Alcian Blue. It is difficult to precisely decipher cell mediated matrix secretion from that already present in the scaffold itself just by staining. The use of non-canonical amino acids described below addresses this limitation. Moreover, we performed biochemical analysis for total GAG content using a DMMB assay, which showed that by day 21 MEND had supported a higher production of GAGs per cell, which was statistically different compared to all other conditions, including pellet controls (Figure 5E).

**Figure 5.**
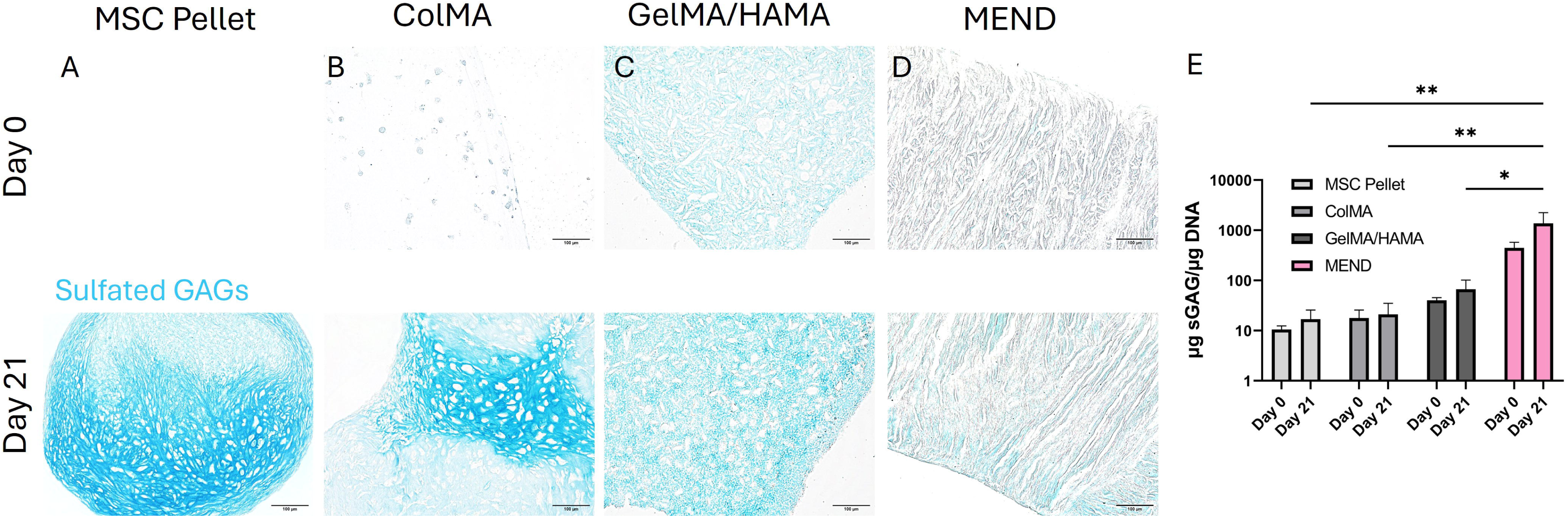
MEND supports uniform matrix secretion and higher GAG production per cell. Alcian blue at pH of 1.0 for **(A)** Day 21 MSC pellet culture, **(B)** Day 0 and Day 21 ColMA, **(C)** Day 0 and Day 21 GelMA/HAMA, and **(D)** Day 0 and Day 21 MEND. Scalebar = 100 µm; all images were taken at 10X magnification. **(E)** Quantification of these results using GAG DMMB biochemical analysis confirms that MEND has a higher GAG production per cell compared to other conditions. Two-way ANOVA, *p<0.05, **p<0.01

### 3.5 MEND allows for MSCs matrix secretion beyond the pericellular space

As many of the components of hydrogels and MEND are also components secreted by cells during chondrogenesis, it may be a challenge to distinguish cell mediated neo-matrix from that of the scaffold itself. To overcome this limitation of biological-based scaffolds, FUNCAT was used to fluorescently label newly secreted matrix. For this, we cultured ColMA, GelMA/HAMA, and MEND constructs in FUNCAT-modified chondrogenic medium where methionine was substituted by homopropargylglycine (HPG), a methionine analog fluorescently taggable by bioorthogonal chemistry. Each sample was sectioned and stained by DAPI (Figure 6A), HPG (Figure 6B), hyaluronic acid binding peptide (Figure 6C), and the overlay of these images (Figure 6D) allowed to observe co-localization of all three channels. Specifically, co-localization of the HA binding peptide and of HPG indicated areas of newly secreted hyaluronic acid. Notably, the HPG signal remained pericellular in both ColMA and GelMA/HAMA conditions, indicative of remodeling limited to the immediate adjacency of cells. On the other hand, MEND constructs exhibited HPG signal further away from the cells, indicating that MSCs in MEND are capable of reaching further in the remodeling of their microenvironment.

**Figure 6.**
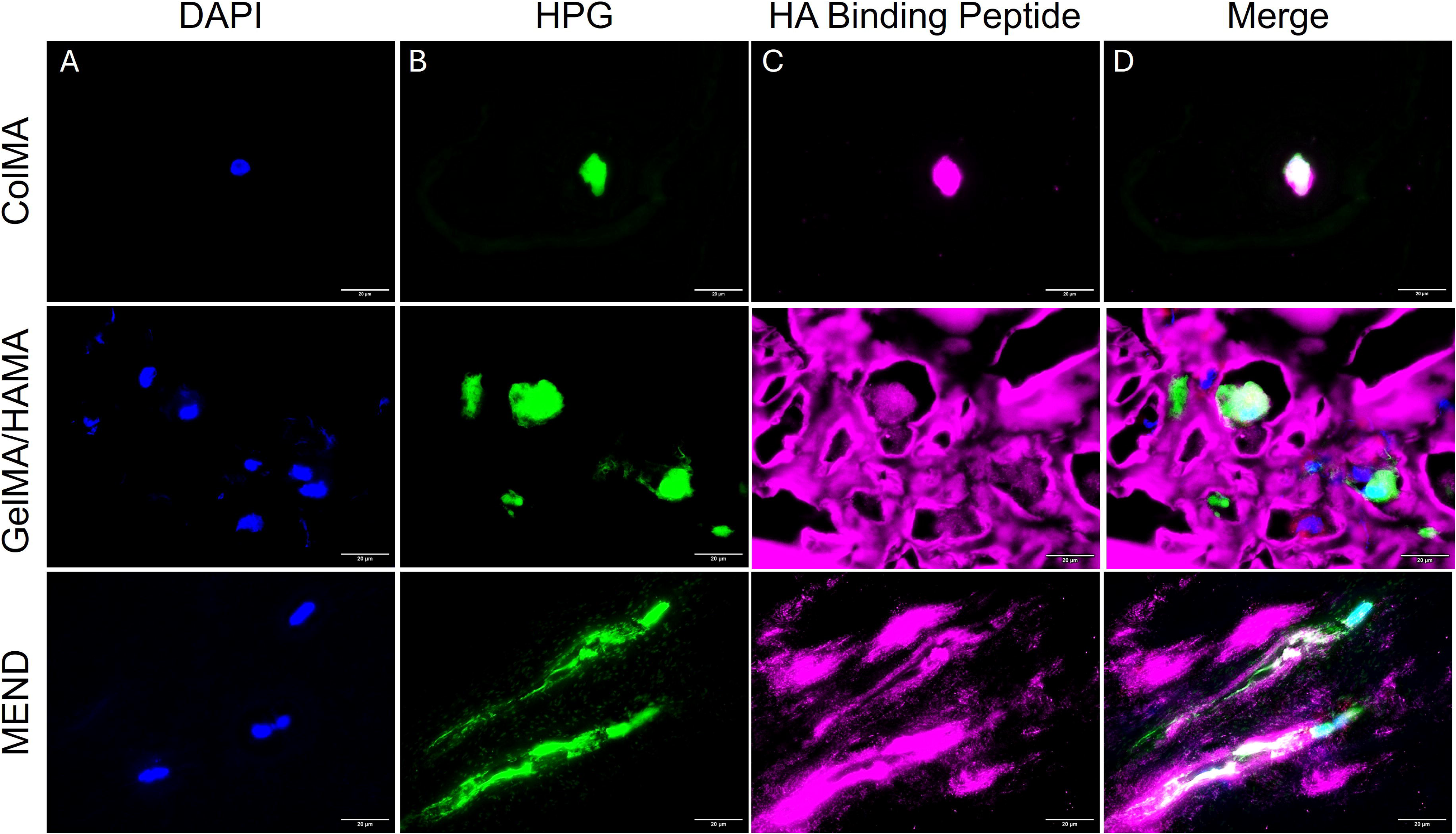
Fluorescent non-canonical amino acid tagging (FUNCAT) highlights that when seeded in MEND, MSCs produce matrix beyond the pericellular space. Samples were cultured in chondrogenic media containing a fluorescently taggable methionine analog for 21 days and were then stained for **(A)** DAPI, **(B)** the methionine analog, homopropargylglycine (HPG), and **(C)** hyaluronic acid binding peptide. **(D)** Stains were merged to reveal that MSCs seeded in all conditions produced neo-matrix containing hyaluronic acid, an important component of cartilage matrix. Scalebar = 20 µm; all images were taken at 100X magnification.

## 4 Discussion

In this work, we assessed the chondroconductive and chondroinductive potential of our dECM scaffold based on porcine meniscus decellularization (MEND). We set out to study MEND’s ability to both support (chondrocondutivity) and promote (chondroinductivity) chondrogenesis in comparison to other tissue engineering materials. To this end, we compared chondrogenesis of MSCs in MEND to that of cells encapsulated in a methacrylated collagen I (ColMA) hydrogel or in a methacrylated gelatin/methacrylated hyaluronic acid (GelMA/HAMA) hydrogel. These hydrogels were selected as ColMA is based on collagen I, representing one of the main ECM components in MEND, and GelMA/HAMA contains hyaluronic acid, another relevant MEND component, thus in principle uncoupling their potential contribution. Our results showed that MEND’s ECM derived composition and novel channel formation allow for chondroconductivity, while chondroinductivity was not substantially enhanced *in vitro* compared to the other tested scaffolds.

After selective enzymatic digestion of meniscal tissue to create MEND, we measured an increase in porosity associated with the removal of elastin, a decrease in DNA content below the FDA approved threshold for decellularization of 50ng of DNA/mg of tissue [28,29], and a decrease in total GAG content compared to native meniscus (Figure 2), similar to what we observed in our previous work [22,23]. We also observed a decrease in compressive modulus which is likely due to the loss of GAGs experienced during the fabrication process. Despite this modulus decrease of about 50%, it is still remarkable that substantive mechanical properties are retained while generating sufficient porosity for uniform recellularization (Figure 3). In fact, complete pulverization of tissue ECM has often been one of the few possible approaches for incorporating decellularized cartilaginous tissues in a tissue engineering approach. This complete loss of native tissue structure drastically reduces mechanical properties to the point of making translation challenging without extensive *in vitro* pre-culture time [30].

In this work, we chose to use MSCs as they are an accessible and potentially autologous stem cell source, increasing their attractiveness as a therapeutic cell type for chondrogenic regeneration [31]. Importantly, MSCs are capable of multilineage differentiation and proliferate faster than chondrocytes [31]. Thus, MSCs provide a solid platform to assess the ability of MEND to support chondrogenesis in comparison to ColMA and GelMA/HAMA.

We seeded MEND with 100k, 400k, and 600k MSCs per scaffold inducing chemotaxis for 5 days to achieve recellularization. We saw that higher cell seeding density resulted in lower cell migration through the scaffold as shown in Figure 3. We hypothesize that this is due to newly formed cell-cell interactions on the surface of the scaffold, which prevents vertical migration through the channels. When a lower density was seeded, cells experienced less cell-cell contact and preferentially travelled towards the nutrient-rich bottom of the transwell, resulting in a uniformly seeded biomaterial as seen in the 100k cells/scaffold condition. Although *in vitro* chondrogenesis routinely requires high cell density to mimic cell condensation and promote cell-cell interactions [32], we seeded cells in a naturally derived ECM environment, requiring a lower cell density that is able to achieve uniform seeding [23].

We then conducted RT-qPCR on all conditions and normalized to day 0 MSC pellets as a positive control (Figure 4). RT-qPCR showed that MEND is comparable to the other materials in supporting upregulation of chondrogenic genes. We did not see strong upregulation of *SOX9* across any of the conditions, but this may be due to our late-stage timepoint. In fact, Lefebvre *et al.* reported that *SOX9* is often expressed in earlier chondrogenesis and its role during late stage chondrogenic differentiation is still not fully defined [33]. Since 21 days is generally considered late-stage chondrogenesis, we expect that this could contribute to our low *SOX9* expression across conditions. Furthermore, RT-qPCR indicated that each of the 3D constructs supported chondrogenic upregulation of *ACAN* in MSCs as compared to donor matched day 0 pellet culture. It is not surprising that higher expression of *ACAN* was observed across conditions compared to *SOX9* since *ACAN* is commonly considered a mature chondrogenesis marker and increases with increasing culture time [34]. Given that a common outcome of many chondrogenic tissue engineering efforts is more similar to fibrocartilage than hyaline cartilage, we also assessed the ratio of *COL2A1*/*COL1A1* to evaluate whether we were achieving more hyaline-like or fibrous-like cartilaginous constructs. We only observed a slight negative trend in the ColMA condition but overall, for all other conditions there was no substantive change in the *COL2A1*/*COL1A1* suggesting that MEND is not better or worse than the other scaffolds in driving a more hyaline cartilage phenotype *in vitro*.

In all constructs, histology via Alcian Blue showed an increase in sulfated GAGs after 21 days of chondrogenic culture, compared to day 0 controls. MEND retained its structure and supported more uniform GAG secretion throughout chondrogenic culture, unlike ColMA and GelMA/HAMA. Although the positive nonspecific hyaluronic acid staining may suggest that GelMA/HAMA supports uniform matrix deposition, the robust mechanical properties and crosslinking modalities of these hydrogels can make them difficult for cells to remodel [35]. To combat this, neo-matrix was identified by including non-canonical amino acids in the culture medium. FUNCAT staining showed that all constructs supported cell mediated neo-matrix secretion. We multiplexed the staining with a hyaluronic acid binding peptide to better understand the amount of hyaluronic acid present within the neo-matrix for each condition. Our results suggest that the MSCs encapsulated in the hydrogels did not manage to substantially remodel the surrounding scaffold, likely because of limited capacity to penetrate a relatively stiff and crosslinked hydrogel. This resulted in a primarily pericellular neo-matrix. Notably, this is a major limitation in the use of GelMA/HAMA as well as other hydrogels. Of note, Xu *et al.* have attempted to overcome this by adding matrix metalloproteinase-cleavable elements in their hydrogels. Such cleavable elements effectively allow for hydrogel degradation over time in response to cell attempts at matrix remodeling, creating more space for matrix secretion and cell migration.[36] Differently from the GelMA/HAMA hydrogel, the ColMA hydrogel contracted substantially during the 21 days culture period. In the presence of cells, we observed the gel starting to shrink within 3 days. In some cases, by day 21 the constructs contracted to the size of a pellet. This is a limitation of many collagen-based hydrogels and the likely reason for the non-uniform matrix secretion in these constructs. Notably, this created regions of higher cell density that may result in GAG non-uniformity throughout the ColMA scaffold [37]. This is a limitation of our work because the high and concentrated expression of Alcian Blue could be a result of contracture, making it difficult to assess GAG secretion produced by MSCs. To combat this, we implemented FUNCAT technology to better decipher cell mediated neo-matrix from that of the construct itself. Notwithstanding this observed hydrogel contraction, FUNCAT staining still highlighted a primarily pericellular deposition of neo-matrix in the ColMA condition. Differently, cells in MEND produced neo-matrix further away from the immediate pericellular space, suggesting that MEND allows for easier matrix secretion compared to other conditions. This is consistent with our previous work where we reported that MEND used *in vivo* for laryngotracheal reconstruction is extensively remodeled [22]. Interestingly, biochemical analysis for sulfated GAGs across all conditions revealed that MEND supports more GAG production per cell. However, despite the higher GAG/cell secretion, our MEND construct appears to be primarily chondroconductive and does not yet show substantive chondroinductivity.

Scaffolds containing dECM have shown chondroinductive properties in many prior studies,[38–40] including some of our own [22]. We previously showed that dECM scaffolds derived from cartilage have enhanced chondroinductive properties compared to hydrogel biomaterials like GelMA [20]. Similarly, Beck *et al.* found that hydrogels composed of devitalized cartilage ECM consistently produced higher collagen II expression compared to hydrogels without cartilage ECM[41]. Our current results do not align with this trend, as MEND appears to be chondroconductive but not chondroinductive. Notably, our work was performed entirely *in vitro* which can often limit the chondroinductive potential of engineered scaffolds. Wang *et al.* created cartilage ECM scaffolds and tested their chondrogenic potential *in vitro* followed by subcutaneous implantation *in vivo*. Their results suggested that *in vivo* implantation led to stronger GAG and collagen production compared to *in vitro* culture [42]. It is then reasonable to speculate that if implanted *in vivo* MEND-based scaffolds might have a higher chondroinductive capacity than we were able to observe *in vitro*. Furthermore, MEND is unique because of its preserved native structure. Creating dECM materials while preserving structure often requires harsh enzymatic treatment which can rid the tissue of intended native cell content but also cause the removal of key signaling components that could contribute to its inherent chondroinductive potential. GAGs such as chondroitin sulfate and aggrecan are often considered important native signaling components to drive chondrogenesis using dECM scaffolds [43]. The use of harsh enzymatic treatment to preserve structure contributed to severe GAG loss during MEND fabrication, which may have led to the low chondroinductivity seen in MEND samples. Ongoing efforts are in progress to improve the fabrication protocol to maximize GAG retention.

Overall, MEND presents important advantages for cartilage tissue engineering compared to other materials currently explored in the field. MEND is uniquely derived from porcine menisci which after decellularization and selective enzymatic digestion results in a cartilaginous matrix biomaterial that retains its structure and integrity. It can be recellularized without requiring pulverization or significant drilling. Hyaline cartilage is known to be extremely dense and difficult to decellularize without inducing damage to the native matrix. MEND decellularization occurs through enzymatic means, which retains matrix properties and decreases toxicity concerns associated with other decellularization protocols involving surfactants. MEND is abundant with channels after the removal of native elastin fibers, opening opportunities for recellularization. Given its natural derivation, cells can easily form interactions with its matrix and migrate through channels, eliminating the need for synthetic manipulations like arginylglycylaspartic acid binding sequences. From a scalability point of view, MEND is a xenogeneic biomaterial which utilizes porcine menisci, a tissue commonly discarded as a biproduct of the food industry, increasing accessibility and decreasing translatability costs. Our results suggest that, although MEND is derived from a fibrocartilaginous source, it does not drive MSC differentiation towards a more fibrous phenotype than ColMA or GelMA/HAMA. MSCs can differentiate equally well in MEND, while presenting superior outcomes in terms of construct remodeling, as seen by the FUNCAT experiments.

This work used human donors to maximize translatability efforts and impact of application for advancing human health. However, human donors result in high levels of donor-to-donor variability. Although we included 5 randomized donors for all outcomes to limit this effect, the high donor-to-donor variability made statistical significance difficult to achieve. This high variability between donors may have also contributed to obfuscating some trends, and it might be a contributing factor in our inability to observe chondroinductive effects of MEND.

MEND introduces promise in providing a biomaterial that is derived from widely available natural tissue, maintains its natural structure and is successfully decellularized. It is also uniformly pervaded by channels which allows for ample recellularization opportunities and supports chondrogenesis to a similar standard in comparison to other tissue engineering materials. MEND has potential to be a complementary biomaterial option for microfracture, a clinical technique largely limited by stem cell clearance due to unavailable attachment sites. MEND could address this limitation by providing a scaffold capable of bone marrow aspirate infiltration to promote stronger chondrogenic regeneration. Additionally, MEND could be applied to other joints that experience osteoarthritis such as the temporomandibular joint.

## ACKNOWLEDGEMENTS

We thank Paul Gehret and Dana D. Ragbirsingh for training on MEND related protocols, Ryan M. Friedman for training on ColMA hydrogel handling, Brennen D. Covely for editing this document, and the Center for Applied Genomic at CHOP for access to their QuantStudio machine.

## FUNDING

This work was supported in part by the NIH/NIAMS T32 Training in Musculoskeletal Research (T32-AR007132) (HMB and KWYS), the Penn Fontaine Fellowship (HMB and KWYS), the Penn Undergraduate Research Mentoring Program (SEK), the CHOP Research Institute Summer Scholars Program for Advancing Child Health: Preparing the Next Generation of Pediatric Researchers (NIH NICHD R25HD101365) (CB), the Penn Center for Musculoskeletal Disorders (NIH/NIAMS P30AR069619), the NIH/NHLBI R56HL164536-01 (RG), the NIH/NHLBI R01HL164536-01A1 (RG), the Children’s Hospital of Philadelphia Research Institute, and the Frontier Program in Airway Disorders of the Children’s Hospital of Philadelphia (RG).

## ETHICAL STATEMENT

No IRB or IACUC approvals were needed for completion of the study.

## CONFLICT OF INTEREST

The authors declare no conflict. RG is co-inventor on a submitted patent application on the meniscus decellularization.

## Notes

### Competing Interest Statement

The authors have declared no competing interest.

