## Supplemental figure legends for "Decellularized Meniscus (MEND) as a biomaterial that supports stem cell invasion and chondrogenesis"

**Supplemental Figure 1.** Alcian blue at pH of 1.0 for human MSC donor 1. **(A)** Day 21 MSC pellet culture, **(B)** Day 0 and Day 21 ColMA, **(C)** Day 0 and Day 21 GelMA/HAMA, and **(D)** Day 0 and Day 21. Scalebar = 100 µm; all images were taken at 10X magnification.

**Supplemental Figure 2.** Alcian blue at pH of 1.0 for human MSC donor 2. **(A)** Day 21 MSC pellet culture, **(B)** Day 0 and Day 21 ColMA, **(C)**Day 0 and Day 21 GelMA/HAMA, and **(D)** Day 0 and Day 21. Scalebar = 100 µm; all images were taken at 10X magnification.

**Supplemental Figure 3.** Alcian blue at pH of 1.0 for human MSC donor 3. **(A)** Day 21 MSC pellet culture, **(B)** Day 0 and Day 21 ColMA, **(C)**Day 0 and Day 21 GelMA/HAMA, and **(D)** Day 0 and Day 21. Scalebar = 100 µm; all images were taken at 10X magnification.

**Supplemental Figure 4.** Alcian blue at pH of 1.0 for human MSC donor 4. **(A)** Day 21 MSC pellet culture, **(B)** Day 0 and Day 21 ColMA, **(C)**Day 0 and Day 21 GelMA/HAMA, and **(D)** Day 0 and Day 21. Scalebar = 100 µm; all images were taken at 10X magnification.

**Supplemental Figure 5.** Alcian blue at pH of 1.0 for human MSC donor 5. **(A)** Day 21 MSC pellet culture, **(B)** Day 0 and Day 21 ColMA, **(C)**Day 0 and Day 21 GelMA/HAMA, and **(D)** Day 0 and Day 21. Scalebar = 100 µm; all images were taken at 10X magnification.
